# Clinically Validated Model Predicts the Effect of Intratumoral Heterogeneity on Overall Survival for Non-Small Cell Lung Cancer (NSCLC) Patients

**DOI:** 10.1101/2021.02.13.431080

**Authors:** Nima Ghaderi, Joseph H. Jung, David J. Odde, Jeffrey Peacock

**Author notes:** Electronic address, Electronic address (David J. Odde, Jeffrey Peacock).

## Abstract

**Purpose:** We demonstrate the importance of considering intratumoral heterogeneity and the development of resistance during fractionated radiotherapy when the same dose of radiation is delivered for all fractions (Fractional Equivalent Dosing FED).

**Materials and Methods:** A mathematical model was developed with the following parameters: a starting population of 10^11^ non-small cell lung cancer (NSCLC) tumor cells, 48-hour doubling time, and cell death per the linear-quadratic (LQ) model with α and β values derived from RSIα/β, in a previously described gene expression based model that estimates α and β. To incorporate both inter- and intratumor radiation sensitivity, RSIα/β output for each patient sample is assumed to represent an average value in a gamma distribution with the bounds set to -50% and +50% of RSIα/b. Therefore, we assume that within a given tumor there are subpopulations that have varying radiation sensitivity parameters that are distinct from other tumor samples with a different mean RSIα/β. A simulation cohort (SC) comprised of 100 lung cancer patients with available RSIα/β (patient specific α and β values) was used to investigate 60Gy in 30 fractions with fractionally equivalent dosing (FED). A separate validation cohort (VC) of 57 lung cancer patients treated with radiation with available local control (LC), overall survival (OS), and tumor gene expression was used to clinically validate the model. Cox regression was used to test for significance to predict clinical outcomes as a continuous variable in multivariate analysis (MVA). Finally, the VC was used to compare FED schedules with various altered fractionation schema utilizing a Kruskal-Wallis test. This was examined using the end points of end of treatment log cell count (LCC) and by a parameter described as mean log kill efficiency (LKE) defined as:

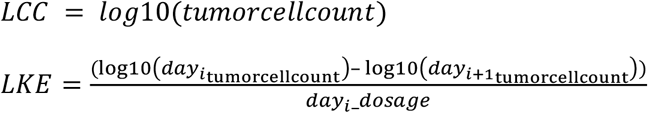

**Results:** Cox regression analysis on LCC for the VC demonstrates that, after incorporation of intratumoral heterogeneity, LCC has a linear correlation with local control (p = 0.002) and overall survival (p =< 0.001). Other suggested treatment schedules labeled as High Intensity Treatment (HIT) with a total 60Gy delivered over 6 weeks have a lower mean LCC and an increased LKE compared to standard of care 60Gy delivered in FED in the VC.

**Conclusion:** We find that LCC is a clinically relevant metric that is correlated with local control and overall survival in NSCLC. We conclude that 60Gy delivered over 6 weeks with altered HIT fractionation leads to an enhancement in tumor control compared to FED when intratumoral heterogeneity is considered.

## Introduction

Radiation therapy is used in nearly 50% of cancer treatments in the developed world (Bodgi et al., 2016). Nearly all radiation treatments today are given over the course of 1-7 weeks with the same dose each treatment day, or fractional equivalent dosing (FED) (Park et al., 2008). The rationale for FED was developed in the 1930s as a method to increase the therapeutic ratio - i.e. ratio of healthy tissue to tumor tissue after radiation - allowing for increased doses to tumor tissue while sparing normal tissue (Fletcher, 1983; Fletcher & Shukovsky, 1975). For the past century, radiation therapy improvements have operated within the constraints of FED. Extensive randomized dose escalation studies across disease sites, apart from prostate cancer, utilizing FED have failed to improve clinical outcomes (Bradley et al., 2015; Jeong et al., 2017; Karim et al., 1996; Minsky et al., 2002; Ramroth et al., 2016; Roach et al., 2018). In previous work, we posited that empiric dosing without taking into consideration intertumoral heterogeneity could explain the lack of impact of dose escalation trials (Hong & Salama, 2016; Scott et al., 2017). However, intratumoral heterogeneity was not considered. An additional explanation for this phenomena is that FED lends itself well to the development of radiation resistance during treatment (Abazeed et al., 2013; Manem et al., 2019; Yard et al., 2016). Intratumoral heterogeneity, highly selective pressure, and rapid turnover are all characteristics of FED that could result in the failure to eliminate the tumor population during radiation treatment (Alfonso & Berk, 2019).

Recent data on exploring intratumor heterogeneity and its impact on fractionated therapy have been explored. The studies that investigate intratumor radiosensitivity utilize the LQ model. The LQ model is the most validated radiation response model for tumor tissue. In this model 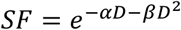, where SF is the survival fraction after radiation while, α (1/Gy) and *β*(1/Gy^2^) are two parameters which map the response of the surviving fraction of a tissue to the quantity of dose applied, D (Gy). However, these works assume an infinite number of non-discrete (continuous) radiosensitivity parameters and tumor proliferation rates that are clinically unrealistic and hence propose solutions that are not clinically feasible, i.e. dose intensities or number of fractions beyond a clinically tolerable limit. Furthermore, models that use the LQ model assume α and β to either be constant over time or have a single value (van Leeuwen et al., 2018).

In addition to theoretical modeling, disease sites that have explored the use of brachytherapy as a boost (i.e. not performing radiation with FED) have been shown to improve clinical outcomes and are now incorporated in standard of care treatments (cervical and prostate cancer). This suggests that the development of resistance to the same fractionated dose developed during radiation and brachytherapy offers a different fractionation modality to overcome the resistance.

To remedy this problem, we propose a new method to incorporate intratumoral heterogeneity through clinically validated estimates of radiosensitivity parameters, providing the platform for a model capable of predicting individual patient outcomes. To clinically validate our model, we focused on NSCLC, which is a leading cause of cancer related mortality in the United States. Roughly 85% of patients have non-small cell lung cancer (NSCLC) and 15% have small cell lung cancer (SCLC). Unfortunately, NSCLC is diagnosed only near advanced staged disease. Treatment is generally resection for stage 1 and 2, or conventional/stereotactic radiotherapy for stages 3 and higher, with or without adjuvant chemotherapy (Duma et al., 2019). Compared to conventional radiation therapy, dose escalation trials have failed to meet their endpoint goal of enhancing overall survival while hypofractionation treatments have been as beneficial if not more efficacious in maintaining or enhancing clinical outcomes compared to conventional radiation treatment in NSCLC (Duma et al., 2019; Hong & Salama, 2016; Jeong et al., 2017; Ramroth et al., 2016; Roach et al., 2018; Scott et al., 2020). In this study, we sought to examine the impact of FED on the development of resistance during radiation as well as possible fractionation schemas to overcome the resistance.

## Methods

### Genomic Methods

MCC Cohorts: The breast and lung cancer cohorts were extracted from Total Cancer Care (TCC), a prospective IRB-approved data and tissue collection protocol active at Moffitt and 18 other institutions since 2006. Tumors from patients enrolled in the TCC protocol were profiled on the Affymetrix Hu-RSTA-2a520709 chip (Affymetrix, Santa Clara, CA), which contains approximately 60,000 probe sets representing 25,000 genes. All of the raw cel files were normalized utilizing RMA. All patients in the TCC cohort were consented for the TCC protocol (Scott et al., 2017).

MCC Simulation cohort – 100 randomly chosen patients with NSCLC cancer profiled under the TCC program were chosen. RSIα/β was used to estimate α and β parameters for each tumor.

MCC Validation cohort – A separate cohort of 57 patients with stage III NSCLC treated at Moffitt with post-operative RT (dose range 45 – 70 Gy). The median follow up was 59.5 months. Clinical outcomes for each patient were available including overall survival and local control.

### Model Development

RSIα/β is a model trained on clonogenic assays performed on NCI60 cell lines to provide a genomic expression estimate of the α and β parameters of the LQ model. There are two distinct models, one for α and one for β, that are clinically validated across three independent cohorts, including data from a phase III randomized clinical control trial. In order to incorporate intratumoral heterogeneity in radiation sensitivity parameters, we assume that the RSIα/β parameters for each patient was derived from a gamma distribution with the mean of the gamma distribution situated at the individual values of each patient’s radiosensitivity parameters.

### Clinical Validation

Clinical validation was performed by another author separately from the model development. Each patient within the validation cohorts RSIα/β parameters and radiation prescription were inputted into the model to determine their survival fraction. Cox regression analysis was used to correlate survival fraction to actual patient time to local control and overall survival in the validation cohort. The other covariates include age, nodal status, and chemotherapy. Age and survival fraction were continuous variables and nodal status (n≥2) and chemotherapy use were categorical.

### Parameter Initialization

Radio-sensitivity parameters for each patient were obtained from TCC Moffitt. These parameters were estimated based on (Scott et al., 2017). Patient tumor doubling times were set to be 2 days extracted from NCI60 data for NSCLC cell lines.

To create radio-sensitivity parameter ranges of the linear quadratic model fit (α,β) [Gy^-1^,Gy^-2^], a Gamma Probability Density Function (GPDF) Random Number Generator in Matlab® is used. The GPDF is described by *γ*:

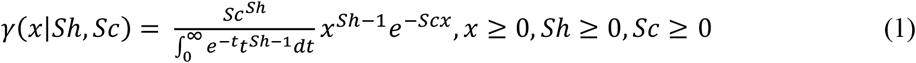

where (Sh, Sc): (shape, scale) parameters ϵ [(0,∞), (0,∞)] and the variable range is x ϵ (0,∞). The mean of GPDF is (Mean= Sh•Sc) and its variance is (Variance=Sh•(Sc^2^)).

The shape/scale ratios of the α distribution and the β distribution were derived from a population of ∼2000 patients. Each patient in the population had individual α and β parameters. The distribution fitter app on MATLAB was used to fit a gamma probability distribution function to the population α data and the population β data, from which the shape/scale ratio was derived. These gamma probability distribution functions were used in all subsequent testing involving the simulation cohort and the validation cohort. The mean of the GPDF is the input patient (α,β) parameter in each case.

In a preoperative scenario, N_0_=10^11^ cells are estimated to be in a tumor. As shown in the supplement, 10 subpopulations (S=10) obtained via draws made from a GPDF to ensure a high probability of at least 1 subpopulation in a tumor that is more resistant to radiotherapy. Each subpopulation is treated as a discrete entity. The initial cell count in each subpopulation is weighed equally, N_i_=N_0_/S_i_. The α and β parameters assigned to each subpopulation are drawn from the α and β gamma distribution derived from the population. These values are limited to a minimum threshold of 50% of the patient’s mean value and a maximum threshold of 150% of the patient’s mean value. If these thresholds are exceeded, draws are rejected and a new draw from the same underlying distribution is undertaken until “S” subpopulations are designated to a patient. The (α,β) are drawn independently from each other. The mean of each parameter is calculated as:

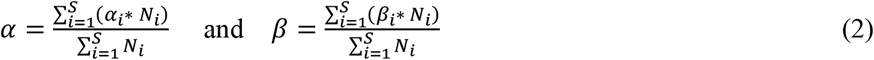

Where each subpopulation is weighted based upon its cell count.

### Treatment Schedule

The radiation therapy schedule is five days a week with two days of rest on weekends, the most common way to deliver fractionated therapy. A given number of fractions (fx) based on clinical recommendations are suggested for a patient, once daily, with a dose intensity of “d” [Gy] / fraction. The dose intensity can be either a constant or variable based on the hypothesis under consideration. In all of the simulated schedules total 60Gy with an average of 2Gy/fraction to be equivalent to 60Gy with 2Gy every fraction (FED). This emphasizes the benefit of altered fractionation without necessitating overall dose escalation.

A total of eight different treatment schedules were studied:

*Conventional Treatments:*

1. The standard of care (FED) is the no-ramp treatment schedule: 30 fractions with 2 Gy/fraction for a total dose delivered of 60 Gy.
2. The ramp-up (RU) treatment schedule increases linearly from 1 Gy on the first treatment day to 3 Gy on the last treatment day. The linear increase occurs exclusively between treatment days (weekdays) and not during days of rest (weekends).
3. The validation treatment schedule was applied during clinical validation of the model. The validation schedule changes per patient and depends on the clinical data which states the number of fractions and daily dose received by each respective patient. *High Intensity Treatments:* The applied proposed dose is beyond 3 Gy, which taps into the quadratic terms of the LQ model based on our estimation of cohort average α and β, is less than the constitutive domain of the validity of the LQ model (dose < 5 Gy), and is still below the 4 Gy clinically tolerable dose (Bodgi et al., 2016). The rationale behind HIT treatments is further explained in the results sections. Briefly, a day of rest is prescribed for a patient after a high dose. The high intensity dose is prescribed to take advantage of the sensitivity of the β parameter to greater dose values. Of note, the total dose remains 60Gy delivered over 6 weeks.
4. High Intensity Treatment_1 (HIT_1): The dosing regimen is: 3 Gy, 1 Gy, 3 Gy, 1 Gy, 2 Gy, 0 Gy, 0 Gy. This regimen repeats for 6 weeks.
5. HIT_2: The dosing regimen is: 4 Gy, 0 Gy, 4 Gy, 0 Gy, 2 Gy, 0 Gy, 0 Gy. This regimen repeats for 6 weeks.
6. HIT_3: The dosing regimen is: 3.33 Gy, 0 Gy, 3.33 Gy, 0 Gy, 3.33 Gy, 0 Gy, 0 Gy. This regimen repeats for 6 weeks.

### Simulation Algorithm

After parameter initialization, time “t” in the simulation is advanced by “Δt” which is dictated by the periodicity of the scheduling (Δt=1 for once daily doses). At each time point “n,” t_n_=t_n-1_ + Δt.

(α,β) is updated based on previous step’s output, with each subpopulation having a weight equal to its cell count for mean estimations. If N_i_<1, cell count in that subpopulation is eliminated.

On rest days, tumor cell population grows exponentially such that, if “gr” is growth rate [1/day], we can write:

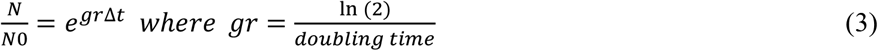

However, if a treatment day is scheduled, upon receiving a fraction of radiation with dose “d,” cell death is assumed to take place first: N_1,2_=N_0_•SF. Survival Fraction (SF) is the ratio of surviving cells to initial cell count after irradiation, given by:

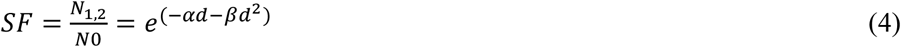

Repopulation is followed such that the final cell count after a day of treatment is N=N_1,2_•e^grΔt^.

In total, the cell count after a fraction of radiation is:

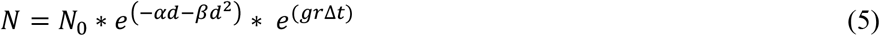

This process is repeated for all subpopulations within a time step, and their cell count is calculated. If (N<1), that subpopulation is assumed to be eradicated. All subpopulations, so long as their count is at least one cell (N≥1), will undergo this selection and elimination process until all time steps in the simulation are completed.

Nine trials are attempted to simulate each patient, with the subpopulation parameters drawn anew from the gamma distributions in each trial. This number of trials is sufficient to guarantee consistency between simulations (p < 0.05). The median across the 9 trials is assigned as the representative value of a single patient. The mean of the representative values across all patients are compared across treatment schedules to compare efficacy. A number of metrics are recorded to display tumor response: α, β, α/β, Log Kill Efficiency (LKE), and Log Cell Count (LCC). LKE and LCC are defined below

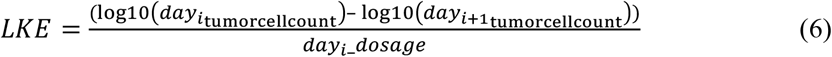

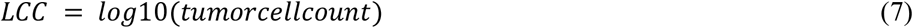

## Results

The model was clinically validated using 57 patients known as the validation cohort. These results are shown in Table 2. The known α, β, and treatment schedules were used as input into the simulation and our model LCC was significant for local control (HR 1.34; p=0.002) and overall survival (HR 1.32; p<0.001) as a continuous variable (Table 2). Hazard ratio > 1.0 signify a better clinical outcome as Age and LCC decrease. Other significant variables include age as a predictor for overall survival. Chemotherapy use trends towards increased local control (p=0.063).

**Table 1:** Description of the various treatment schedules. Treatment 3 is not represented in the table as individual doses vary per patient.

| Treatment Schedule | Monday | Tuesday | Wednesday | Thursday | Friday | Saturday | Sunday | Mean Dose |
| --- | --- | --- | --- | --- | --- | --- | --- | --- |
| (1) FED | 2 | 2 | 2 | 2 | 2 | 0 | 0 | 2 |
| (2)* RU | 1 | 1.0690 | 1.1379 | 1.2069 | 1.2759 | 0 | 0 | 2 |
| (4) HIT-1 | 3 | 1 | 3 | 1 | 2 | 0 | 0 | 2 |
| (5) HIT-2 | 4 | 0 | 4 | 0 | 2 | 0 | 0 | 2 |
| (6) HIT-3 | 3.33 | 0 | 3.33 | 0 | 3.33 | 0 | 0 | 2 |

\*continues to rise linearly by 0.0690 Gy until the final day of treatment, where the final dosage is equal to 3 Gy

**Table 2:** p-value of Cox regression analysis of validation cohort.

|  | Local Control |  | Overall Survival |  |
| --- | --- | --- | --- | --- |
|  | p-value | 95% CI | p-value | 95% CI |
| **LCC** | 0.002 | 1.34 (1.11 – 1.56) | $<0.001$ | 1.32 (1.13 – 1.52) |
| **Age** | 0.736 | 1.00 (0.966-1.050) | 0.005 | 1.06 (1.02-1.10) |
| **Chemotherapy** | 0.063 | 7.21 (0.934-13.48) | 0.964 | 1.35 (0.46-1.52) |

**Table 3:** The “Difference of Last Day LCC” is the representative LCC value of FED on the last day (day 42) subtracted by the representative LCC value of RU, HIT_1, HIT_2, or HIT_3 on the last day. The “Mean Difference of LKE” is the mean of the representative LKE values across all days of FED subtracted by the mean of the representative LKE values across all days of RU, HIT_1, HIT_2, or HIT_3.

| FED vs. | RU | HIT_1 | HIT_2 | HIT_3 |
| --- | --- | --- | --- | --- |
| **Difference of Mean LastDayLCC (FED - X)** | 0.28 | 0.56 | 1.65 | 1.52 |
| **Kruskal-Wallis p-value (LastDayLCC)** | 0.51 | 0.23 | p < 0.001 | p < 0.001 |
| **Difference of Mean LKE (FED - X)** | -0.0005 | 0.016 | -0.037 | -0.046 |
| **Kruskal-Wallis p-value (LKE)** | 0.37 | 0.31 | 0.23 | p < 0.001 |

We performed simulations on 100 patients known as the simulation cohort and obtained their α, β, α/β, LCC, and LKE trends over time (Figures 1-3). Two scenarios of RU and FED were implemented on the simulation cohort. Analysis of the FED condition in Figure 1(a)-1(c) revealed that selective pressures result in the emergence of resistance subpopulations within approximately 10 days of treatment. From this, it was hypothesized that by implementing a RU treatment schedule, the patient would benefit as the higher doses of radiation toward the end of the treatment regimen would be more effective against the resistant subpopulations. However, as seen in the simulation of RU conditions (Figure 2), the effects on final cell count and LKE were marginal. Further mathematical investigation, as highlighted in the supplemental section, revealed that for the clinically relevant α and β parameters of NSCLC, the final cell count depends only on the average dose intensity and not on the treatment schedule unless the applied dosing schedule involves high intensity treatment as explained below.

**Figure 1:**
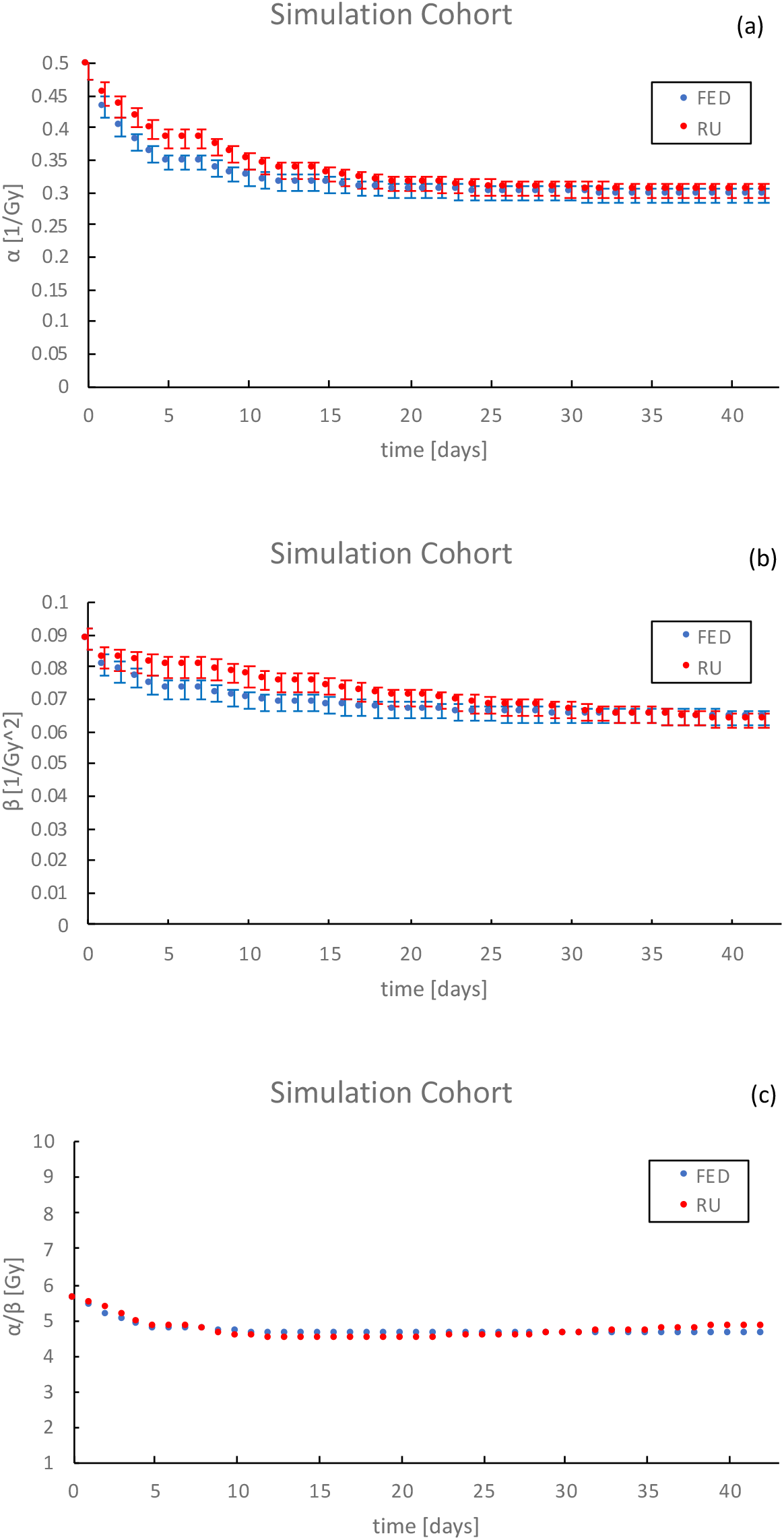
Parameter trends for the simulation cohort. (a) Average simulation cohort α are shown for standard of care (FED) and ramp up (RU). The emergence of resistance is demonstrated at ∼ day 15 before which α drops significantly but after which it remains approximately constant. Error bars are SEM. (b) The mean β for the simulation cohort as a function of time, error bars are SEM. (c) The mean α/β values as a function of time. As α/β drops in time, patients will receive potential benefit from dose escalation. The error bars represent the entire range of patients rather than actual variations in the α/β ratio.

**Figure 2:**
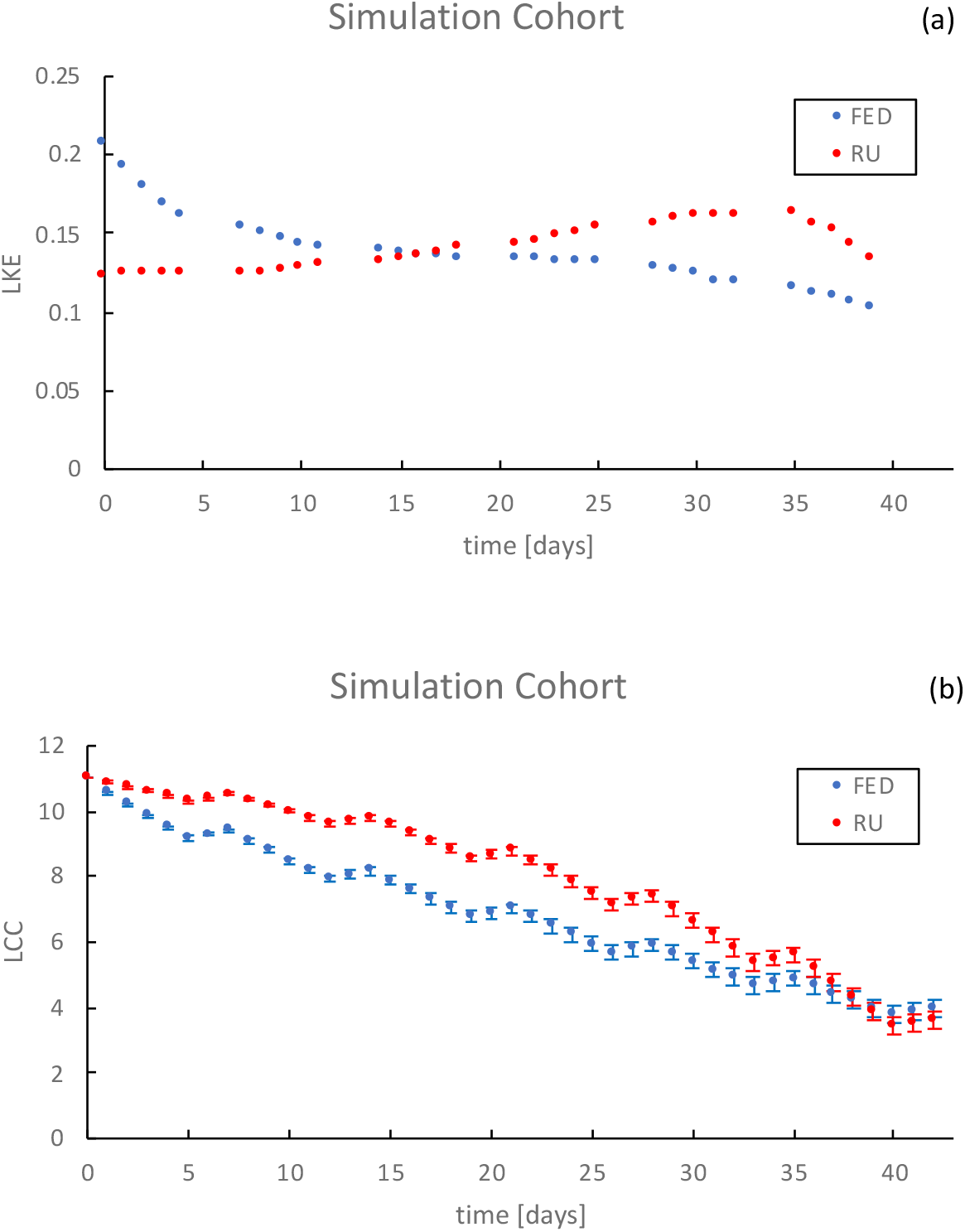
Simulated radiation efficiency (LKE) and tumor size (LCC) as a function of time. (a) Efficiency of the radiation dosage over time. The error bars represent the entire range of patients rather than actual variations in the α/β ratio. (b) Tumor over time. Despite consistent radiation efficiency for RU, the p-value of Kruskal-Wallis analysis between LCC of the last treatment day is not significant (p=0.31). Error bars are SEM.

**Figure 3:**
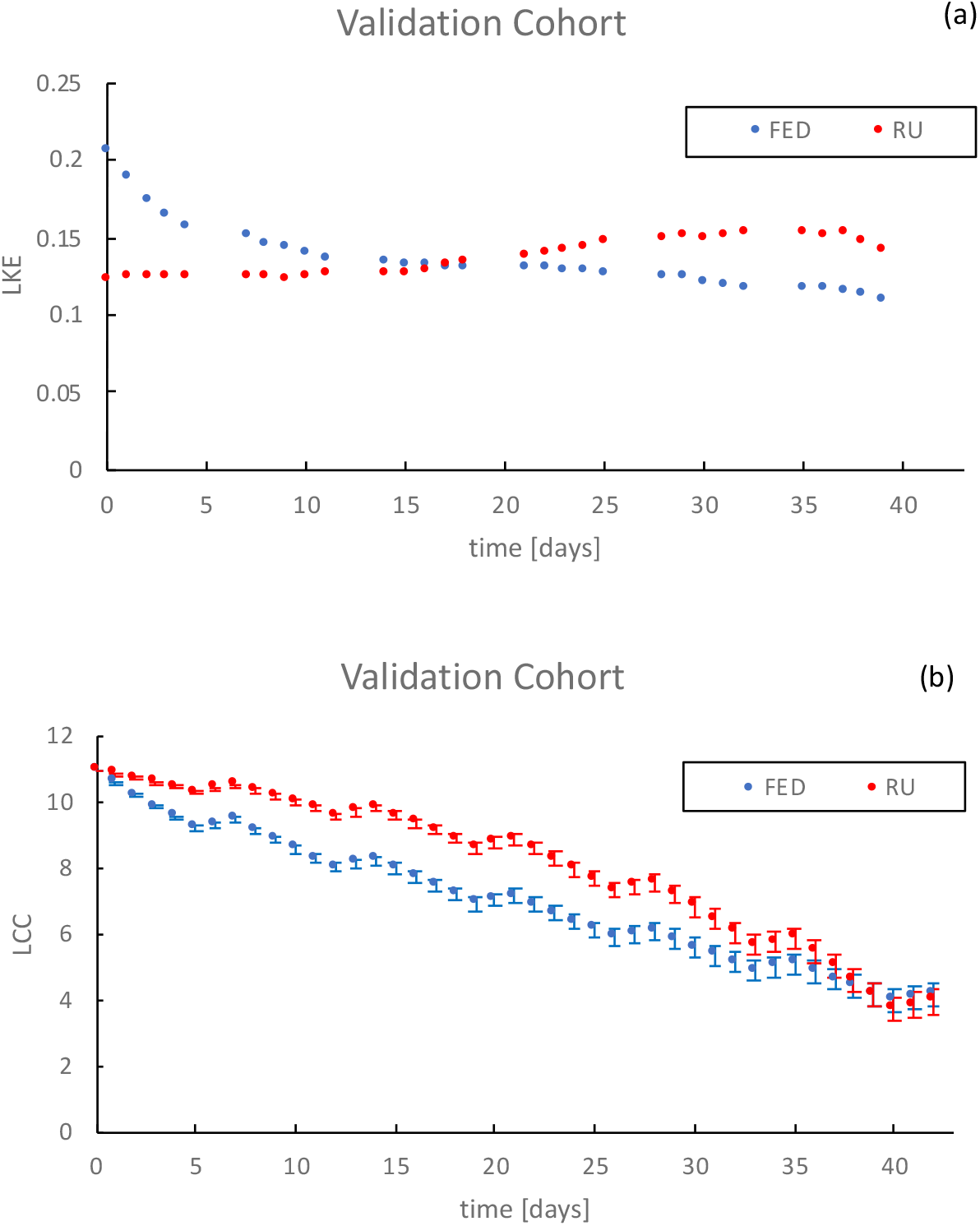
Output trends in the validation cohort of radiation efficiency (LKE) and tumor size (LCC). (a) Efficiency of the radiation dosage over time. The error bars represent the entire range of patients rather than actual variations in the α/β ratio. The general decrease of LKE is due to the emergence of resistant subpopulations. (b) Size of the tumor over time. If average dose is constant, there is no significant benefit in tumor control (FED, RU). See Table 2 for p-values. Error bars are SEM.

As Figures 1(a)-1(c) suggest, the efficiency of the radiation, or LogKillEfficiency, decreases in the FED case as the resistant subpopulations persist. However, in the RU case, the LKE increases as the remaining subpopulations are more susceptible to high degrees of radiation. Based on this conclusion, we hypothesized that by increasing the intensities of dose while maintaining a constant average dose, there would be a higher efficiency in eradication of the tumor (Figure 4). Figure 4(a) shows that LKE is consistently higher for HIT treatments, supporting our conclusion that HIT dosing schedules are tied to favorable outcomes. In Figure 4(b), there is a significant drop in final cell count, with HIT_2 appearing the most effective, followed by HIT_3. HIT_1 was not found to have any significant improvement in the simulated outcomes. These schedules are constrained by current clinical standards of care limitations; however, we understand that they are not definitively the most efficient scheduling methods for treatment.

**Figure 4:**
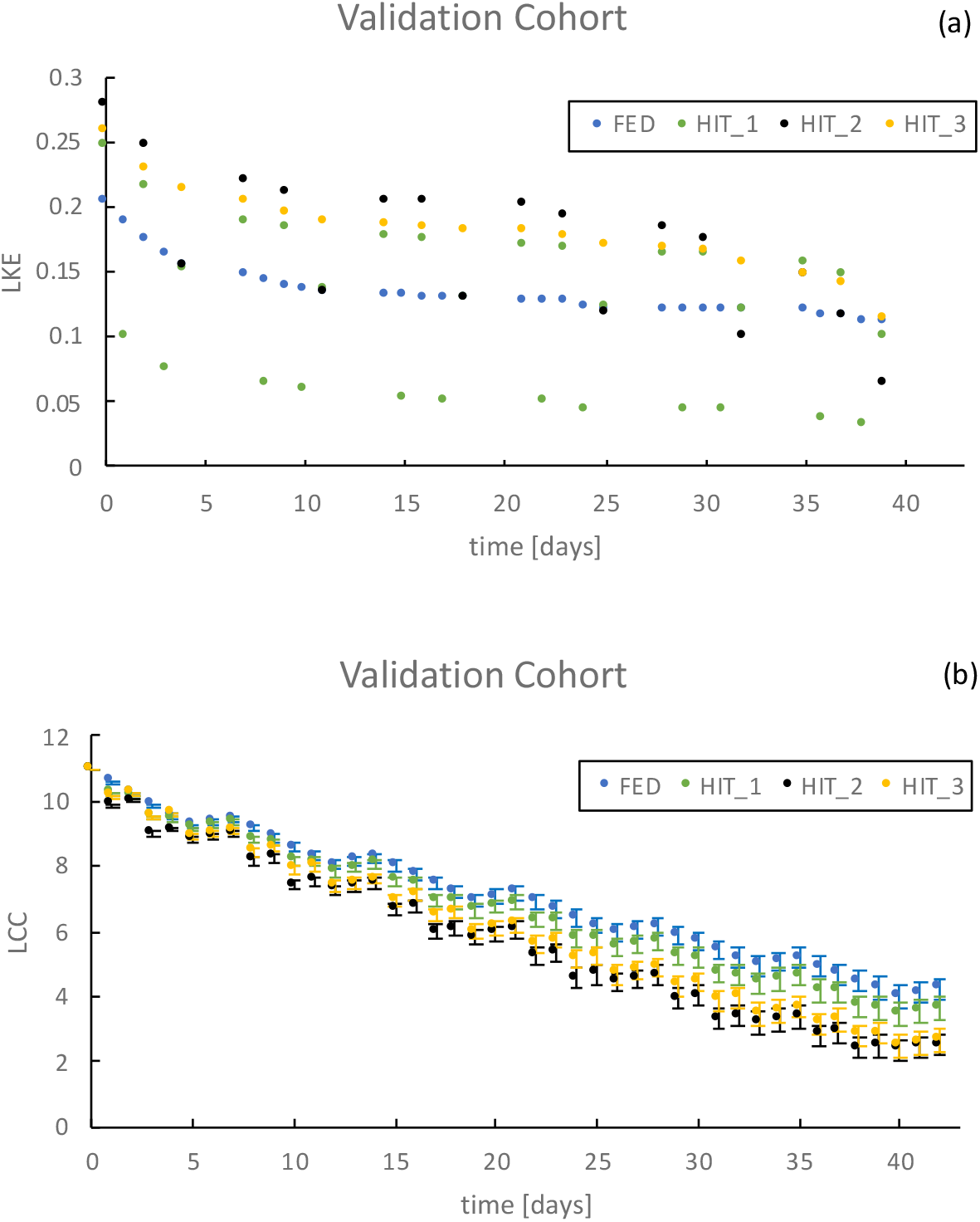
Output trends in the validation cohort of radiation efficiency (LKE) and tumor size (LCC). (a) Demonstrates the efficiency of the radiation dosage over time. Error bars were not included as they represent the range of patients in the simulation cohort and not actual variations in LKE. (b) Demonstrates the size of the tumor over time. All HIT treatments demonstrate a benefit in tumor control, with HIT_2 and HIT_3 being the most effective. See Table 2 for p-values. Error bars are SEM.

As is shown in figure 3(b), RU results in small and insignificant, improvements in the eradication of tumor for individual patients. This was informed by the emergence of resistance towards the end of treatment. Additionally, given the average parameter values for α and β parameters, we demonstrated that patients will receive benefits if higher dose per fraction, still within clinically feasible ranges, are administered to patients. It should be noted that not all patients would require the same schedule treatments as the differences between patients and the heterogeneity within the patient can be determinants of the personalized medicine required for the maximum benefit.

## Discussion

The underwhelming results of radiotherapy dose escalation trials in lung cancer and most disease sites (except for prostate cancer) could be attributed to the development of resistance during radiation when delivered in FED. Despite our knowledge of the development of resistance towards other treatment modalities including systemic therapy and antibiotics, nearly all radiation treatment today continues to be performed using FED, a concept nearly a century old (Martin, 1935).

In our previous work, we demonstrated that interpatient radiosensitivity heterogeneity can result in vastly different clinical outcomes when radiation is delivered in a “one size fits all” approach (Scott et al., 2020). This model seeks to capture intratumor heterogeneity by utilizing a gamma distribution of RSIα/β. This is a reasonable assumption given that RSIα/β was trained in a histological agnostic approach where biological heterogeneity in radiosensitivity will manifest itself in the α and β parameters. Through this clinically validated model, we show that patients will on average have better therapeutic outcomes if the current standard of care is modulated to account for the emergence of resistance within the subpopulations of each tumor. Namely, by keeping the average and total dose constant, we demonstrate that the heavy intensity treatment schedules HIT-2 and HIT-3 will theoretically yield significantly better results. With LCC being a clinically validated predictor for local control and overall survival in FED, we posit that HIT regimens would offer superior clinical outcomes from the theoretical decrease in LCC.

This key finding is further supported by our mathematical formulation provided in the supplement. By assuming patient-to-patient variation in radiosensitivity parameters and heterogeneity within a patient, we were able to demonstrate that regardless of the α and β parameters within the course of a radiation treatment for a cohort of patients, the ratio of the quadratic term to the linear term in the survival fraction will remain substantially below 1 (∼0.2) for the vast majority of fractions. This will result in the LQ model to be approximated with a linear trend, rendering dose escalation treatments ineffectual as the overall effect would be independent of dose scheduling and rather depend on average dosing. Our findings are consistent with RTOG0617, a phase III randomized trial that compared 60Gy in 30 fractions vs. 74Gy in 37 fractions show no improvement in local control or overall survival.

The clinically validated model endpoint, LCC, was shown in the supplement materials to be dependent on the average dose and not individual fraction values so long as the applied fractions are less than 3 Gy. This limit was demonstrated in Table S1. Therefore, in order to improve clinical outcomes, we demonstrated that an increase in dose intensity would shift the ratio of the quadratic portion to the linear portion of the survival fraction to values significantly close to 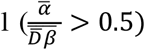 with 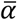 and 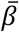 being the mean cohort radiosensitivity parameters and 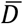 being the average dose applied in a radiation schema. This dimensionless number defines the threshold at which the ratio of the quadratic term and linear term in the LQ model becomes prominent. This will in turn elicit a greater response from the quadratic portion and hence reduce the survival fraction of the tumor, causing the more tumor cytotoxicity. This was demonstrated by the simulations of the heavy-intensity treatments (HIT). Hence, we were able to identify a potential reason behind failure of dose escalation clinical trials in eliciting a response and provide mathematical and simulation proof that hypofractionation can be more beneficial for enhancing overall survival than FED or dose escalation.

### Limitations

- vascularization and damage thereto are not considered
- subpopulations do not evolve during the course of radiation
- concurrent chemotherapy was not modeled in this mathematical formulation
- spatial heterogeneity is not accounted for
- patient performance factors, previous drugs, chemotherapy are not considered
- assumption of RSIα/β interpatient deviation as an approximation for intratumor deviation in RSIα/β
- the simulation endpoint of survival fraction is a surrogate for the effectiveness of treatment, thus complete tumor eradication is not necessary for a patient to not reoccur
- intra-patient α and β values are bound to 50% and 150% of the average value of each respective patient

## Supplemental

### Subpopulation Number Choice

The number of subpopulations (S) chosen in a scenario for a patient reflects the degree of intratumoral heterogeneity; i.e. S=1 represents no heterogeneity (i.e. homogeneous) and S=10 represents the heterogenous case. This is different than the time evolution of individual (α,β) parameters within the tumor. The pool of (α,β) chosen initially to represent intratumoral heterogeneity remains invariable with time, as suggested by clinical data (MOFFITT). But the selection pressure of the irradiation is such that the initial distribution of these parameters vary as more radiosensitive subpopulations die out with initial fractions.

Subpopulation number choice affects the consistency of the outcome for a simulation. If (α,β) are independent variables, randomly chosen from the pool of parameters, the likelihood of having an adverse set of α and β for an individual subpopulation “i” augments as the number of draws (subpopulation count) increases. In other words, for large enough of S, it is highly likely that one pair of (α,β) will be highly radioresistant. Since, in theory, having at least one or more resistant clones negatively affects the likelihood of success of the radiation therapy, we could write down the probability of having one or more resistant clones in the initial subpopulation based on a Binomial distribution. This value signifies successfully drawing “i” adverse pairs with success chance of “p” and failure chance of “q=1-p” out of “S” subpopulations.

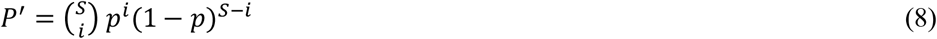

Having at least one or more adverse pairs results in one or more resistant clones, so the actual probability of having embedded resistance due to primary (α,β) parameter initialization is the sum of the probability of having 1,2,3,…,S resistant clones out of S draws.

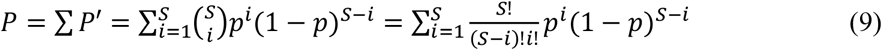

Based on Stirling’s approximation for large factorials, it can be shown that:

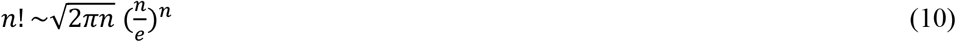

Therefore, the factorial part in (9) can be re-written as:

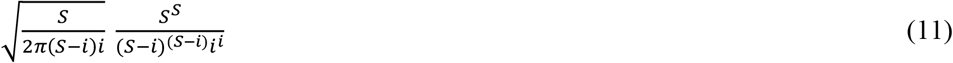

Which yields:

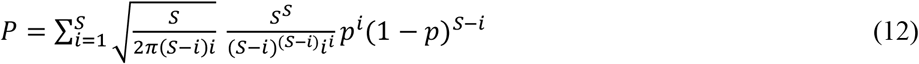

The choice of defining the appropriate “p” value (probability of finding a pair of (α,β) which is resistant) can be formulated as follows.

Based on the LQ model, very low α and very low β are adverse to radiation response. If the entire range of α and β are assumed to be divided into two equal ranges of low-mid-high values, respectively, then the probability of choosing two independent variables at random, for both α AND β to be in their low ranges, is p = 1/2 • 1/2 = 1/4.

As demonstrated, if the inherent probability (p) of having a low chance of drawing adverse combinations of (α,β) is high, number of draws subpopulations will unlikely affect the drawing of at least one adverse combination. However, if (p) is low, the more draws are taken, the more likely it will be that after (i) draws, an adverse combination will have been selected out of the pool of possible (α,β) parameters. Probability of p = 0.25 and S = 10 will almost guarantee that there will be at least one inherently resistant subpopulation drawn.

### Dosing and Growth Rate

Tumor growth rate can be modeled using the following mathematical expression:

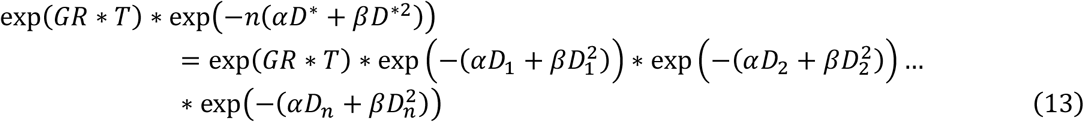

Where GR is growth rate, T is time in days, D is dose in Gy, n is the total number of treatment days, D_n_ is the treatment dose at day n in Gy, 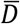 is the average dose across n days in Gy, and D* is a single dosage representative of the dosage across n days in Gy.

Eliminating the growth rate term present on both sides:

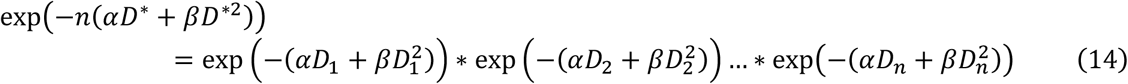

Combining the exponential terms:

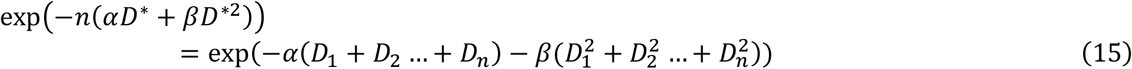

Eliminating the exponential function present on both sides and dividing by n:

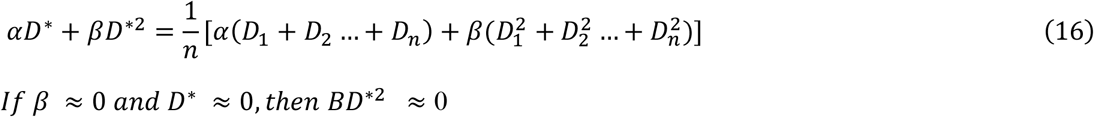

*If β* ≈ 0 *and D*\* ≈ 0, *then BD*\*^2^≈ 0

Assuming small β values:

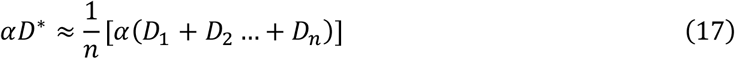

And dividing by *α* yields:

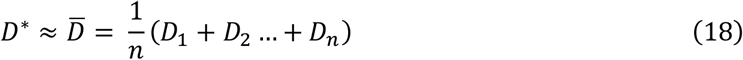

Thus:

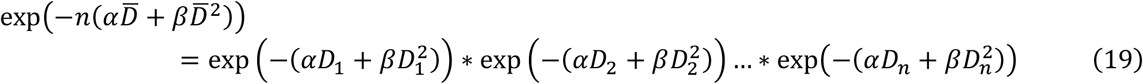

Yields the conclusion that growth rate is independent of *D*_1_,*D*_2_,…*D*_n_ and depends only on 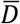.

### Cohort Mean Analysis

Average cell count of all patients can be expressed as:

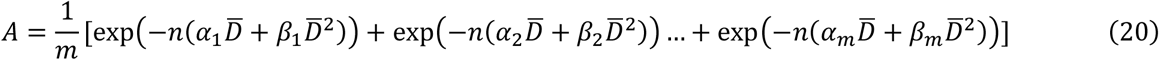

Where A is the average cell count of all patients, D is the dose in Gy, *n* is the total number of treatment days, *m* is the total number of patients, 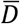 is the average dose across *n* days, is the α value of patient *m*, and is the value of patient *m*.

Multiplying A by m:

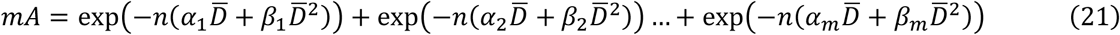

Making a first-order Taylor series approximation for the exponential function:

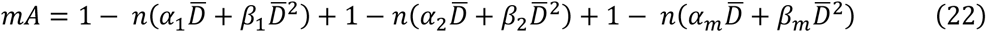

Simplifying the summation of ones to m:

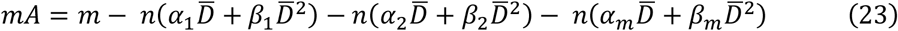

Dividing by m and rewriting in summation notation:

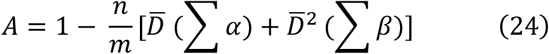

And employing the definition of average to yield:

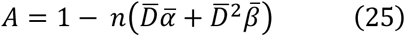

Which shows that the average cell count of all patients at the end of the treatment is independent of individual α, β values and depends only on their averages 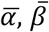 as well as the average dose 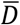, provided that individual doses are below 3 Gy. This is the mathematical threshold at which, given 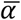 and 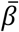, the ratio of 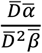 becomes prominently close to 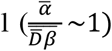. This is a novel dimensionless number that we believe represents the threshold of significance for the quadratic portion of the LQ model. Our results show that better outcomes via HIT-2 or HIT-3 schedules can be achieved theoretically if this dimensionless number is below unity.

Cohort mean α and β values at the start and end of a conventional radiation treatment are (α_*start*_ = 0.496, (β_*start*_ = 0.0886, (α/β_*start*_ = 5.599) and (α_*end*_0.294, β_*end*_ = 0.0643, α/β_*end*_ = 4.573) respectively. The figures below demonstrate why, after a drop in α/β, increasing the dose enhances the ratio of the quadratic term to the linear term in the survival fraction of the LQ model. For doses below the constitutive domain (D < 5 Gy), proposed dosing beyond 3 Gy increases the influence of the quadratic term and makes the treatment more effective theoretically. The reason behind our proposed limit of 4 Gy was informed by current clinical practices, but if safety could be established above this limit, our results predict better outcomes even with no change in total dose.

**Table S1:** The reciprocal of the dimensionless number 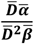 for various doses.

| $((D^2 \beta) / D \alpha)$ | D = 1 Gy | D = 2 Gy | D = 3 Gy | D = 4 Gy | D = 5 Gy |
| --- | --- | --- | --- | --- | --- |
| Start | 0.1786 | 0.3573 | 0.5359 | 0.7145 | 0.8931 |
| End | 0.2187 | 0.4374 | 0.6561 | 0.8748 | 1.0935 |

### Distribution of Subpopulation Parameters

**Figure S1:**
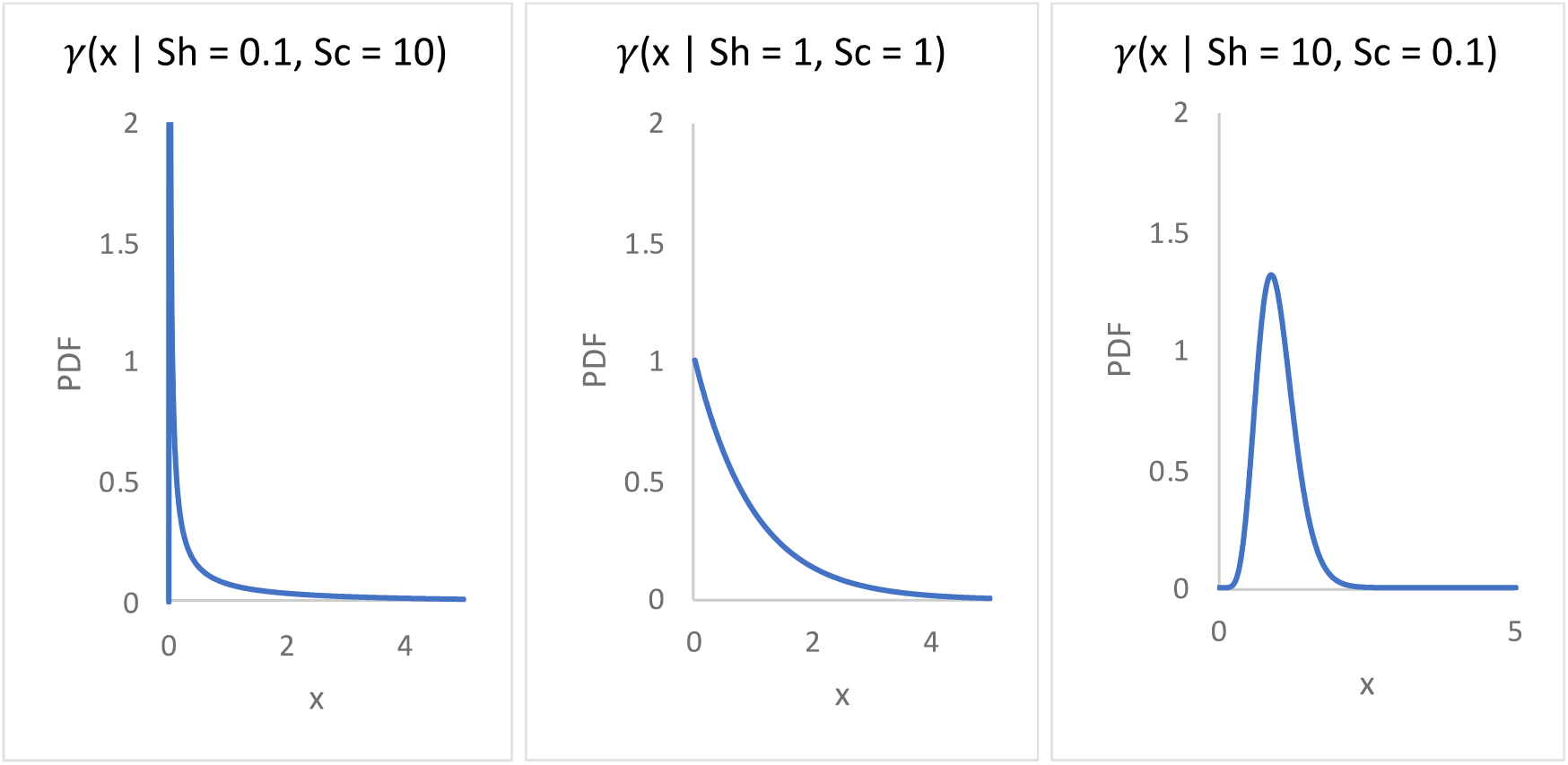
Probability density functions (PDF) of gamma distributions with constant mean (Shape • Scale = 1) but varying Shape/Scale ratios.

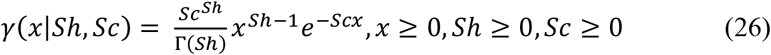

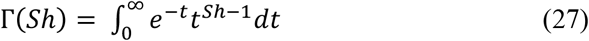

**Figure S2:**
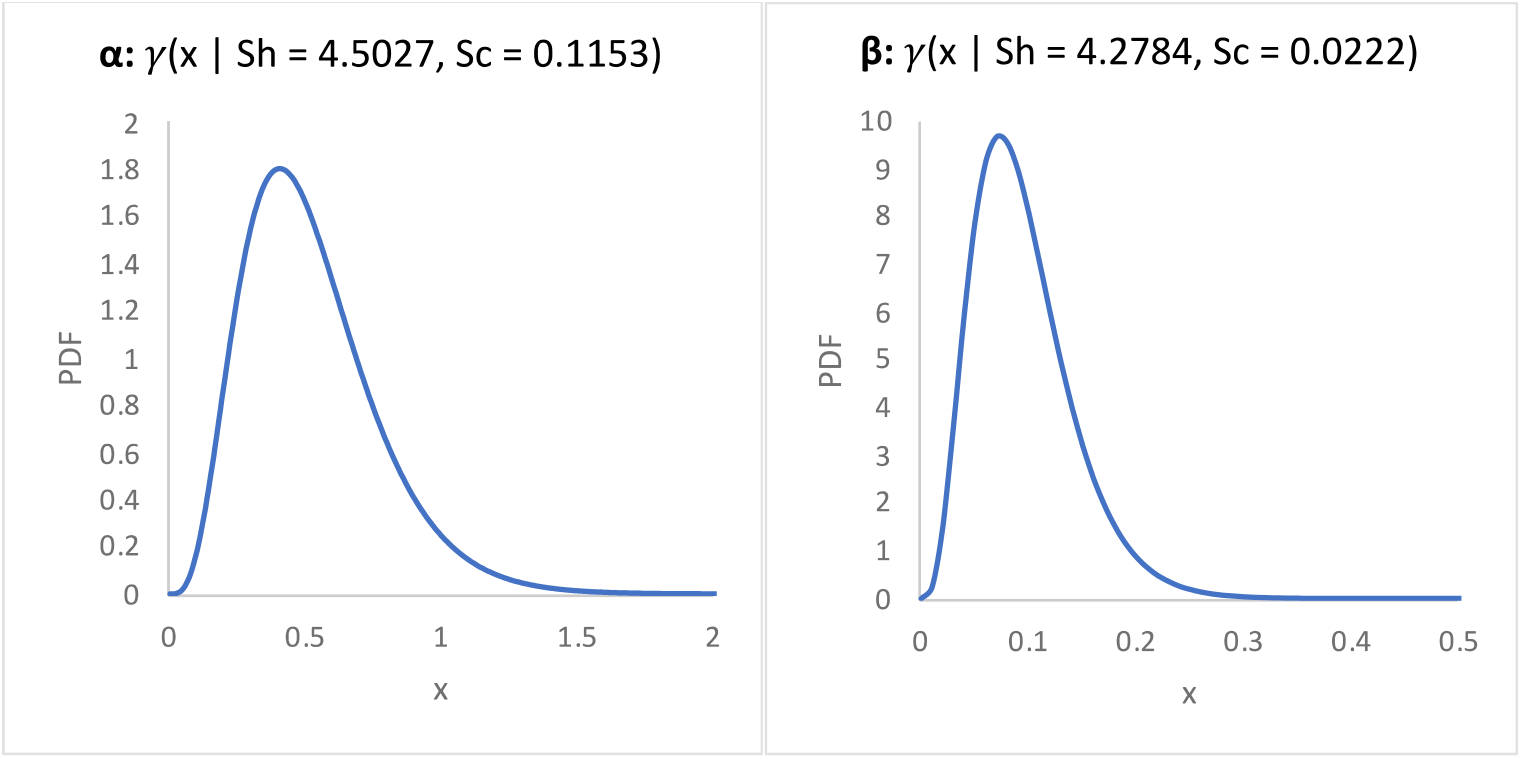
Population probability density functions (PDF) of α [1/Gy] and β [1/Gy^2^].

**Figure S3:**
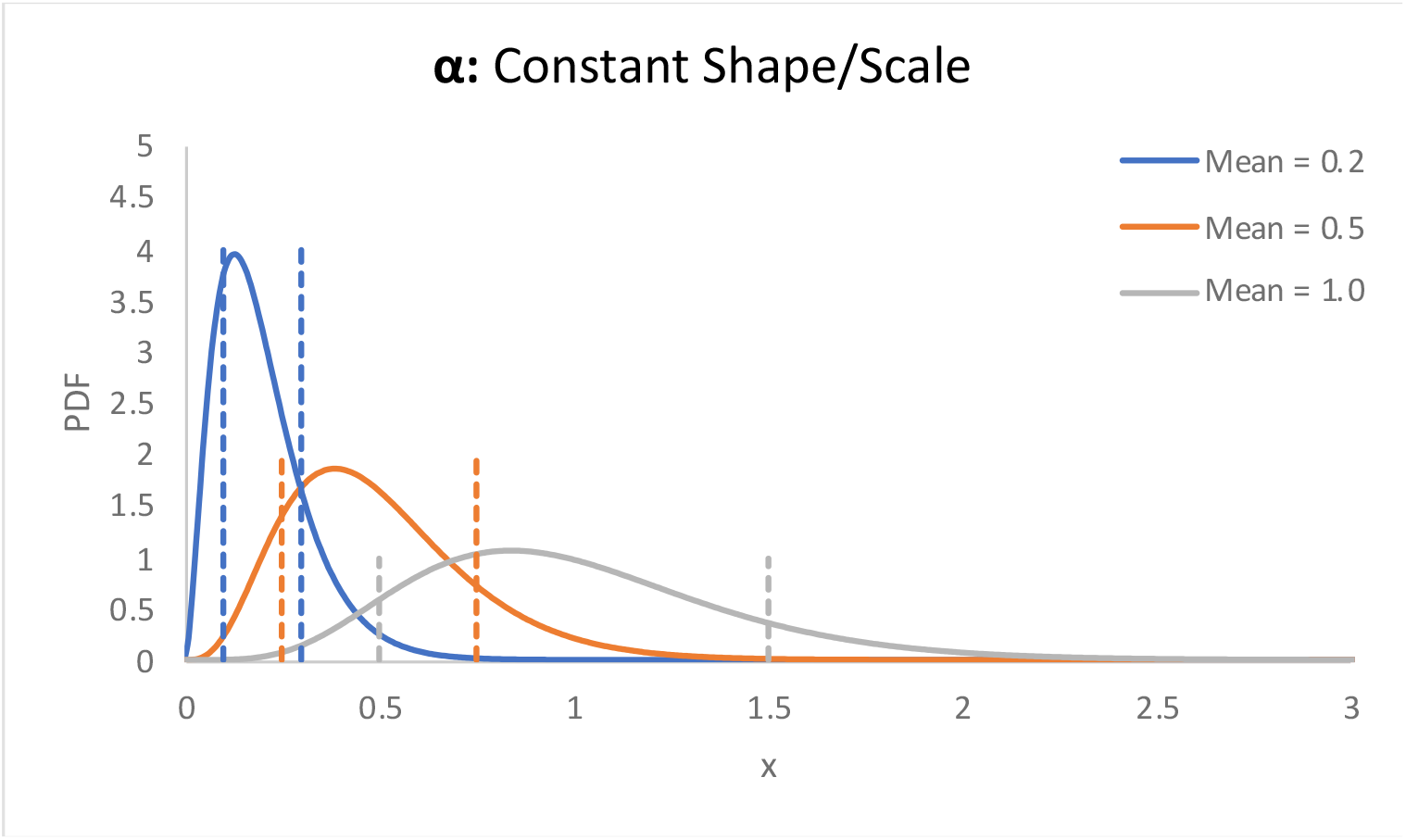
Probability density function (PDF) of gamma distributions with constant Shape/Scale ratios but varying means (Shape • Scale).

Figure S1 demonstrates the different shapes of the gamma distribution with a mean (Sh • Sc) of 1 and a shape to scale ratio (Sh / Sc) of 0.01, 1, and 100 from left to right. In order to maintain a similarity between different patient’s distributions, it was decided to maintain the shape to scale ratio as a constant.

Figure S2 demonstrates the α and β distributions of the entire population cohort, and Figure S3 demonstrates 3 representative α values of 0.2, 0.5, and 1.0 with their respective gamma probability density functions and ±50% mean cutoff points.

